# In vitro neuroendocrine differentiation of BPH-1 cells under β2 adrenergic receptor stimulation

**DOI:** 10.1101/2024.12.04.626774

**Authors:** Natascha Pigat, Laura Nalleli Garrido Castillo, Coline Lefevre, Manon Baures, Vincent Goffin, Julien Anract, Anne-Sophie Armand, Thierry Capiod

**Affiliations:** Inserm U1151, Institut Necker Enfants Malades (INEM), Paris, France; Hôpital Cochin, Paris, France

## Abstract

Epidemiologic and experimental animal studies have shown that stress can alter tumor growth and that adrenergic fibers from the sympathetic nervous system, acting through stromal β2 adrenergic receptors, promote tumor cell survival during the initial phases of cancer development. Recent epidemiological data suggest that beta blocker intake improves survival of prostate cancer patients. Hypertension and high blood pressure have been suggested to increase likelihood of prostate cancer development, suggesting that the use of beta blockers in these patients could reduce prostate cancer risk. Signaling mediated by β adrenergic receptors promotes neuroendocrine differentiation in the LNCaP prostate cancer cell line. Although neuroendocrine cells are a minor population in the epithelial compartment of normal prostate glands, the population of neuroendocrine-like cells that express markers such as SYN, Bcl-2, Dll-3, and AMACR correlates with cancer progression, androgen-independent state, and poor prognosis in prostate cancer patients. As recent studies show that an increase in neuroendocrine cell density occurs before development of benign prostatic hyperplasia, a non-lethal disease affecting the transition zone, in hypertensive rats, we investigated whether neuroendocrine differentiation is associated with benign prostatic hyperplasia. Our results showed that isobutyl-methyl-xanthine, which blocks cAMP degradation, and forskolin, an adenylate cyclase activator, triggered sustained activation of the cAMP/PKA pathway and led to an increase in neuroendocrine differentiation markers in BPH-1 cells, which are a benign prostatic hyperplasia cell line. Similar results were observed by combining the β adrenergic receptors agonist isoproterenol with forskolin. The same treatments reduced BPH-1 cell proliferation rates by 40% and 60%, respectively. Effects on AMACR and SYN expression and cell proliferation were fully reversed by the β2 blocker carvedilol. These data suggest that β adrenergic receptors are linked to the development of neuroendocrine-like cells and benign prostatic hyperplasia. As 20% of the prostate cancer tumors occur in this zone, this observation warrants further investigation.

## Introduction

Stress can influence tumor growth as shown in several epidemiologic and experimental animal studies (1-3); however, the biological mechanisms underlying such effects are not well understood, and the clinical significance of stress on progression of human disease remains controversial. Recent studies indicate that adrenergic fibers from the sympathetic nervous system, acting through stromal β2 and β3 adrenergic receptors, promote tumor cell survival during the initial phases of prostate cancer development (4-6). Innervation patterns of human prostate cancer support this hypothesis: Sympathetic fibers accumulate in normal tissues and infiltrate the tumor edge (7). Data from animal models suggest that stress modulates the growth of certain tumors via neuroendocrine regulation of the immune response to tumor cells (1).

Beta blockers, which are competitive agonists of the ligands of β adrenergic receptors, are among the most commonly used medications in the treatment of hypertension, although recent meta-analyses have concluded that β2 blockers are inappropriate first-line agents in the treatment of hypertension (8). Recent epidemiological data indicate that β2 blocker intake is associated with improved survival of prostate cancer patients (9, 10) and that hypertension and high blood pressure can increase prostate cancer development (11, 12), suggesting that the use of beta blockers could benefit to these patients.

The β adrenergic receptors signal through the cAMP/PKA pathway, a pathway known to induce neuroendocrine differentiation (NED) in LNCaP prostate cancer cell lines (13, 14). Neuroendocrine (NE) cells are a minor cell population in the epithelial compartment of normal prostate glands; the population of NE-like cells is higher in prostate tumor lesions and correlates with tumor progression, poor prognosis, and the androgen-independent state (15). NE-like cells have higher levels of tumoral marker AMACR, SYN, and Bcl-2 which is indicative of resistance to apoptosis, than NE cells (15). DLL-3 regulating Notch signaling (16) is another marker of NE cells in several tissues, including prostate, (17-20) which expression is increased following PKA pathway activation (21).

Recent studies have shown that an increase in NE cell density occurs before development of benign prostate hyperplasia (BPH) in spontaneously hypertensive rats (22). BPH is a non-lethal disease affecting the prostatic transition zone where adenomatous nodules mainly appear. NE cells are distributed around small adenomas and influence the initial growth of BPH in human patients (23). Interestingly, about 20% of all prostate cancers originate in transition zone (24). Furthermore, AMACR expression is detected within evolving carcinomas in some BPH nodules and in fully developed grade 1 carcinomas (25). In most cases, prostatic biopsies used to screen for prostate cancer target the peripheral zone, where most cancers occur, and transition zone cancer is often detected during BPH surgery (10% incidence among all BPH surgeries) (26).

We speculated that β2 adrenergic receptors influence the development of NE-like cells in BPH and that β2 blockers may reduce NED in this zone. We therefore investigated the relationship between NED and BPH in vitro. BPH-1 prostate cells were challenged with a β adrenergic receptor agonist isoproterenol (ISO) or with the combination of isobutyl-methyl-xanthine (IBMX) and forskolin (FSK) to stimulate the cAMP/PKA pathway. NE-like cell markers were analyzed to detect differentiation. By RT-qPCR and western blot, we showed that these treatments resulted in the expression of NED markers in these cells. Furthermore, we showed that β2 blockers antagonized the effects of β2 adrenergic receptor agonists.

## Materials and methods

### Cell culture

BPH-1 (benign prostate hyperplasia), RWPE-1 (normal epithelia), LNCaP (lymph node metastasis), DU-145 (brain metastasis), and PC-3 (bone metastasis) cells were purchased from ATCC (Manassas, USA) and used before passage 20. They were maintained in DMEM with Glutamax-I supplemented with 200,000 U/ml penicillin and 100 μg/ml streptomycin (Gibco, Les Ulis, France), and with 10% fetal calf serum (Eurobio Scientific, Les Ulis, France).

### Cell proliferation, cytotoxicity, and viability

For proliferation, BPH-1 (400 cells/well), PC-3 (1200 cells/well), DU-145 (1200 cells/well), LNCaP (2000 cells/well), and RWPE-1 (1200 cells/well) cells were seeded in 96-well plates. After 3 days of culture, for all cells except the BPH-1 cells, the medium was replaced with the same culture medium containing the test compounds and Red NucLight (1/1000) for staining of nuclei. All drugs were added simultaneously in our experimental conditions. Plates were transferred to an Incucyte plate reader (Sartorius, Dourdan, France) for an additional 4 days. The Incucyte software allowed detection and numeration of red nuclei every hour over this 4-day period. Results given here are the values at the end of the 4-day period of acquisition. For BPH-1 cells, cell proliferation was assessed by measuring the cell confluence using the same software. BPH-1 confluence was always less than 80% at the end of the experiments. BPH-1 cell viability was assessed using Trypan Blue staining quantified on a Biorad TC 20 (Bio-Rad, Marnes-la-Coquette, France) after 4 days of treatment with appropriate drugs.

### Western blot

Fresh cells were pelleted, then lysed for 30 min at 4 °C in RIPA buffer. After centrifugation, the supernatant was evaluated for protein quantity using the Pierce BCA KIT (Sigma-Aldrich, Saint-Quentin-Fallavier, France). Protein samples (30 µg) were electrophoresed in a 4-12% gradient SDS-PAGE in NuPAGE Bis-Tris Precast Gels (Life Technologies, Saint Aubin, France). Proteins were then transferred onto nitrocellulose membranes (Bio-Rad), and membranes were incubated with primary antibodies (Table S1). For band detection, HRP-coupled secondary anti-rabbit (7074, Cell Signaling, Saint-Cyr-L’École, France) or anti-mouse (7076, Cell Signaling) antibodies were added before ECL substrate (Immobilon Western Chemiluminescent HRP Substrate, Millipore, Guyancourt, France).

### qPCR

Total RNA was isolated from cell pellets using the NucleoSpin^®^ RNA XS (Macherey Nagel, Hoerd, France) according to manufacturer’s instructions. RNA (250 ng) was reverse transcribed using PrimeScript™ RT Master Mix #RR036A (Takara Bio Europe SAS, Saint Germain en Laye, France), The cDNAs were then subjected to real-time PCR amplification using gene-specific primers (0.5 µM final concentration) purchased from Sigma-Aldrich (Saint-Quentin-Fallavier, France) (Table S2). *PPIA*, which encodes Cyclophilin A, was analyzed as a control in each reaction. Real-time PCR was performed using a Qtower 2.0 (Analytik Jena, Germany). Each qPCR reaction contained 2 µL cDNA sample (12.5 ng) and 18 µL mastermix with 1X GoTaq^®^ qPCR Master Mix (Promega, Charbonnières les Bains, France) and 0.5 µM primer. The Qtower 2.0 Instrument was used with the following program: enzyme activation: 95 °C for 2 min; amplification (40 cycles): 95 °C for 15 s; and 60 °C for 60 s. Results were generated with the Qtower 2.0 software and were analyzed by the comparative cycle threshold method and presented as fold change in gene expression relative to internal calibrators as mentioned in figures. Experiments were performed in duplicate, and the results are expressed as means ± S.D.

### Statistical analysis

A paired-samples t-test was used for statistical analyses of differences in cell proliferation. As the Incucyte technique allowed high-throughput screening, each set of data was compared to its own control on the same plate and overall differences were tested using the statistical test. As control values differed from one plate to another (Supp. Figure 1A), we normalized the data to controls on each plate. Results are expressed as means ± S.E.M. A one-way ANOVA was used for statistical analysis of western blot data with *, ** and *** corresponding to p values <0.05, <0.01 and <0.001 respectively.

### Compounds

IBMX, FSK, ISO, propranolol, norepinephrine, epinephrine, and phenylephrine were purchased from Sigma-Aldrich, carvedilol and atenolol were purchases from Cayman Chemicals (Interchim, Montluçon, France), mirabegron was purchased from Tocris (BioTechne SA, Noyal Châtillon sur Seiche, France), and NucLight was purchased from Sartorius (Dourdan, France).

## Results

### Stimulation of the cAMP/PKA pathway inhibits proliferation of prostate cancer cells

As the aim of this study was to investigate whether stimulation of the cAMP/PKA pathway could induce NED of BPH-1 cells, we first evaluated whether the combined effects of IBMX and FSK reduced proliferation of these cells; this combination was previously been shown to do so in other cell lines (14). DMSO, which was used as a vehicle, did not affect the rate of cell proliferation compared to distilled water. The mean proliferation rates corresponded to a 12.0 ± 1.0 and 12.0 ± 0.9 fold increase from initial values (n=30) without and with 0.02% DMSO, respectively (Supp. Figure 1A, left panel), corresponding to 3 to 4 cell cycles over a 96-h period of time, with a range in proliferation rate of approximately 4 to 28 fold increase in number of cells treated or not with DMSO (Supp. Figure 1A, right panel). Normalization to cells not cultured with DMSO showed a range from 0.8 to 1.2 units (Supp. Figure 1B), and drug effects were therefore compared to proliferation rates in their respective control conditions throughout this study. The combined effects of 100 µM IBMX and 10 µM FSK reduced cell proliferation by 40%, whereas IBMX alone no effect and FSK reduced proliferation by about 20% (Figure 1A). The combination of IBMX and FSK also reduced proliferation of RWPE-1, LNCaP, DU-145, and PC-3 cells, other prostate lines, with comparable efficiency (Supp. Figure 1C). These results indicate that blocking cAMP degradation with IBMX was less effective than adenylate cyclase activation with FSK in halting cell proliferation but that the combination was more effective than either drug alone. Importantly, the combination had no effect on cell viability (Supp. Figure 2).

**Figure 1:**
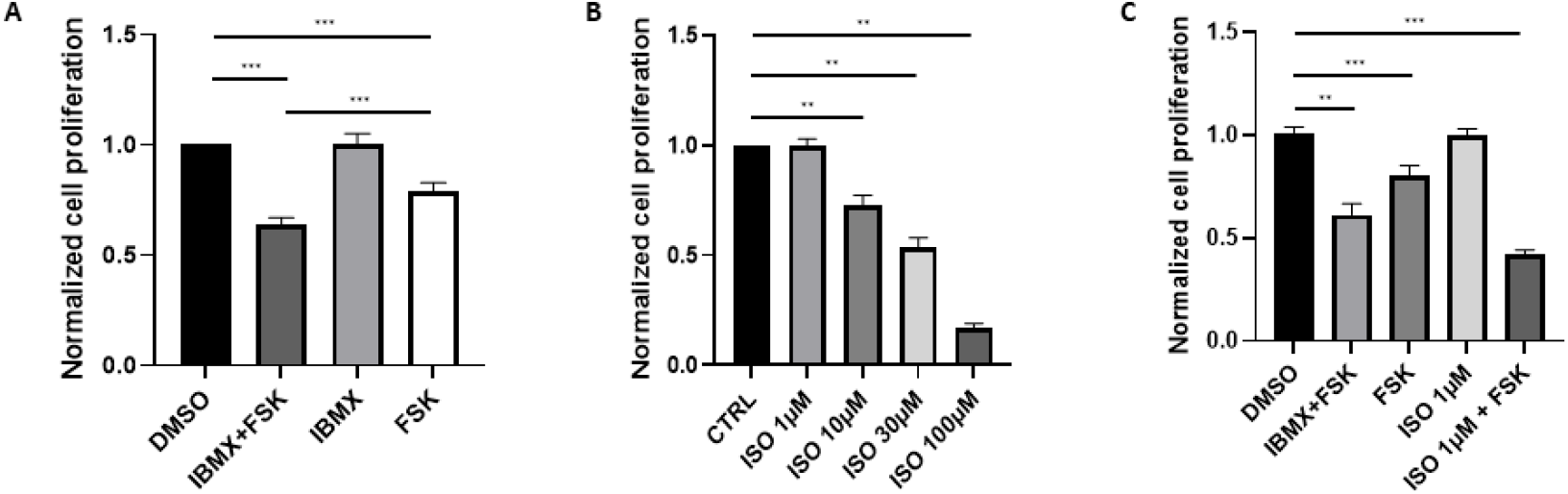
Agonists of β-adrenergic receptors inhibit proliferation of BPH-1 cells. Cell proliferation was measured after 96h in BPH-1 cells treated with (A) 100 µM IBMX and 10 µM FSK (n=23), FSK alone (n=29), or IBMX alone (n=19); (B) indicated concentrations of ISO (n=8-24); and (C) the combination of 100 µM IBMX and 10 µM FSK, 10 µM FSK alone, 1 µM ISO alone, and the combination of 1 µM ISO and 10 µM FSK (n=5-22). DMSO is the vehicle (n=30). Values were normalized to untreated cells, i.e. distilled water for ISO and DMSO for all other conditions.

### Agonists of β adrenergic receptors and beta blockers reduce BPH-1 cell proliferation

IBMX and FSK induce cAMP production in prostate cell lines as well as ISO (14, 27, 28). Therefore, we tested the effect of ISO on BPH-1 cell proliferation. ISO treatment reduced BPH-1 cell proliferation in a dose-dependent manner (Figure 1B). The large decrease in cell proliferation observed at 100 µM ISO was associated with a decrease in cell viability, but lower concentrations of ISO did not kill cells (Supp. Figure 2). The ISO-induced decrease in BHP-1 cell proliferation was also observed in other prostate cell lines (Supp. Figure 3). We next tested the combined effects of a low concentration of ISO and FSK. ISO at 1 µM, a concentration that did not inhibit cell proliferation, combined with 10 µM FSK induced a 60% reduction in cell proliferation, a larger decrease than that observed in the presence of the combination of 100 µM IBMX and 10 µM FSK (Figure 1C).

Norepinephrine and epinephrine are natural neurotransmitters released from nerve terminals that do not discriminate between α and β adrenergic receptors. Both norepinephrine and epinephrine inhibited BPH-1 proliferation (Supp. Figure 4). As the α1 adrenergic receptors agonist phenylephrine had no effect on cell proliferation (Supp. Figure 4), norepinephrine and epinephrine appear to act exclusively through β2 adrenergic receptors in the prostate cancer cells. Mirabegron, a β3 adrenergic receptor agonist (29), caused a small reduction in BPH-1 cell proliferation at 30 µM (Supp. Figure 4). Some studies have shown that *ADRB3* is expressed in prostate tissue (30), but we cannot rule out an effect of mirabegron through β2 adrenergic receptors at this concentration, and this will need further investigation. In summary, these experiments showed here that reduction in BPH-1 cell proliferation is associated with β2 adrenergic receptor activation.

Next we tested whether beta blockers inhibit cell proliferation independently of β adrenergic receptor stimulation by ISO. To do so, we used two different β2 blockers and one β1 blocker. Atenolol, a β1 blocker, had no effect on BPH-1 cell proliferation at concentrations up to 100 µM, but both β2 blockers, propranolol and carvedilol, reduced cell proliferation at 20 and 5 µM, respectively (Figure 2). Increasing the concentration of these two β2 blockers further inhibited cell proliferation via a mechanism that could involve mitochondrial bioenergetics as suggested based on studies of medulloblastoma (31). Tests in three other prostate cell lines confirmed our observations (Supp. Figure 5). Thus, β2 blockers affect prostate cancer cell proliferation independently of β2 adrenergic receptor activation as shown previously in gastric adenocarcinoma, colon adenocarcinoma, and non-small cell lung cancer (31). However, the mechanisms of action of these blockers in BPH-1 cells and other prostate cell lines remain unknown.

**Figure 2:**
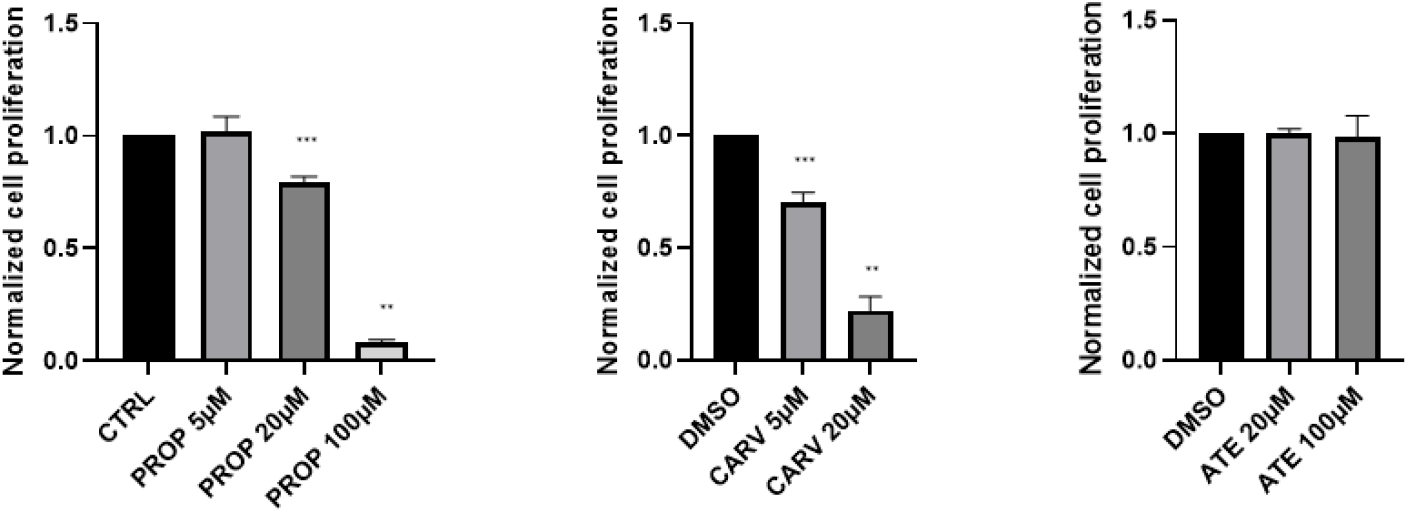
Beta blockers reduce BPH-1 cell proliferation. Normalized BPH-1 cell proliferation was measured after 96h in the presence of propranolol (PROP) (n=6-27), carvedilol (CARV) (n=5-18), and atenolol (ATE) (n=4-17). Normalization was done versus distilled water for PROP or DMSO for beta blockers.

Next, we tested whether these beta blockers could reverse the inhibitory effect of the combination of ISO and FSK on proliferation of BPH-1 cells. As shown before, 1 µM ISO combined with 10 µM FSK induced a 60% decrease in cell proliferation. This was totally reversed in the presence of 5 µM carvedilol and almost completely reversed in the presence of 20 µM propranolol (Figure 3). Atenolol had no effect on cell proliferation inducted by ISO and FSK (Figure 3). This confirmed that β2 adrenergic receptors are responsible for inhibition of proliferation of BHP-1 cells.

**Figure 3:**
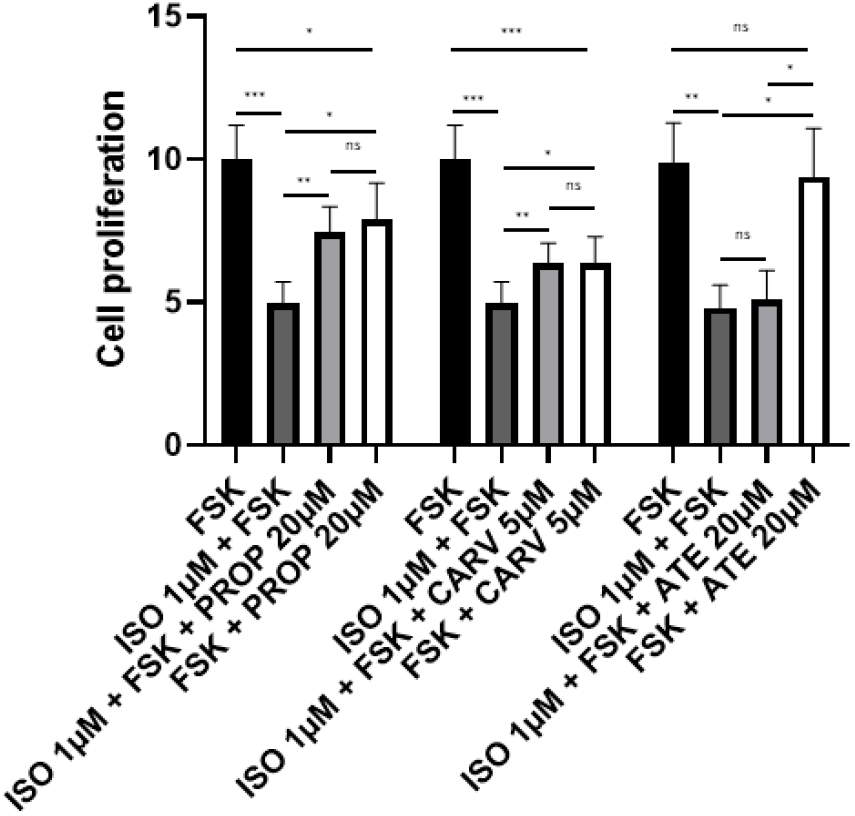
Beta blockers inhibit the effects of β-adrenergic receptor agonist isoproterenol. Cell proliferation was assessed after 96 h in the presence of 10 µM FSK or the combination of 1 µM ISO and 10 µM FSK with or without 20 µM propranolol (PROP), 5 µM carvedilol (CARV), or 20 µM atenolol (ATE) (n=7-8).

### Activation of β adrenergic receptors induces neuroendocrine differentiation of prostate cancer cells

Markers of NED include SYN, Dll-3, Bcl-2, and AMACR (15, 20). We therefore evaluated how activation of β adrenergic receptors and beta blockers affected the expression of these proteins. BPH-1 cells were treated for 10 days with the combination of IBMX and FSK or the combination of ISO and FSK in the presence or absence of β2 blockers. SYN, Dll-3, and Bcl-2 proteins were significantly increased in the presence of the combination of IBMX and FSK and in the presence of the combination of ISO and FSK (Figure 4 and Supp Figure 6). The β2 blockers 20 µM propranolol and 5 µM carvedilol both decreased the effects of the ISO and FSK combination with a clearer effect for carvedilol, the more active blocker of cell proliferation, than propranolol. Another feature of NED differentiation is AMACR expression. AMACR was barely detectable in BHP-1 cells; it is expressed at much higher levels in LNCaP cells (Figure 4D). By western blot, AMACR levels in BHP-1 cells were increased by treatment with IBMX and FSK and with ISO and FSK, but differences were not significant in the presence and absence of carvedilol. qPCR experiments, however, showed that the mRNAs encoding AMACR and SYN were increased in the presence of the combination of IBMX and FSK and by the combination of ISO and FSK and that the latter effect was reversed by 5 µM carvedilol (Figure 5A and B). Interestingly, *SYN* mRNA expression was not increased in RWPE-1 cells under IBMX and FSK treatment, suggesting that NED does not occur in normal prostate cells under these conditions (Figure 5C). *SYN* mRNA expression in RWPE-1 was about 90% lower than in BPH-1 cells (Figure 5C). Overall, these data showed that BPH-1 cells are more susceptible than normal non-tumoral cells such as RWPE-1 to differentiation into NE-like cells upon β2 adrenergic receptor stimulation.

**Figure 4:**
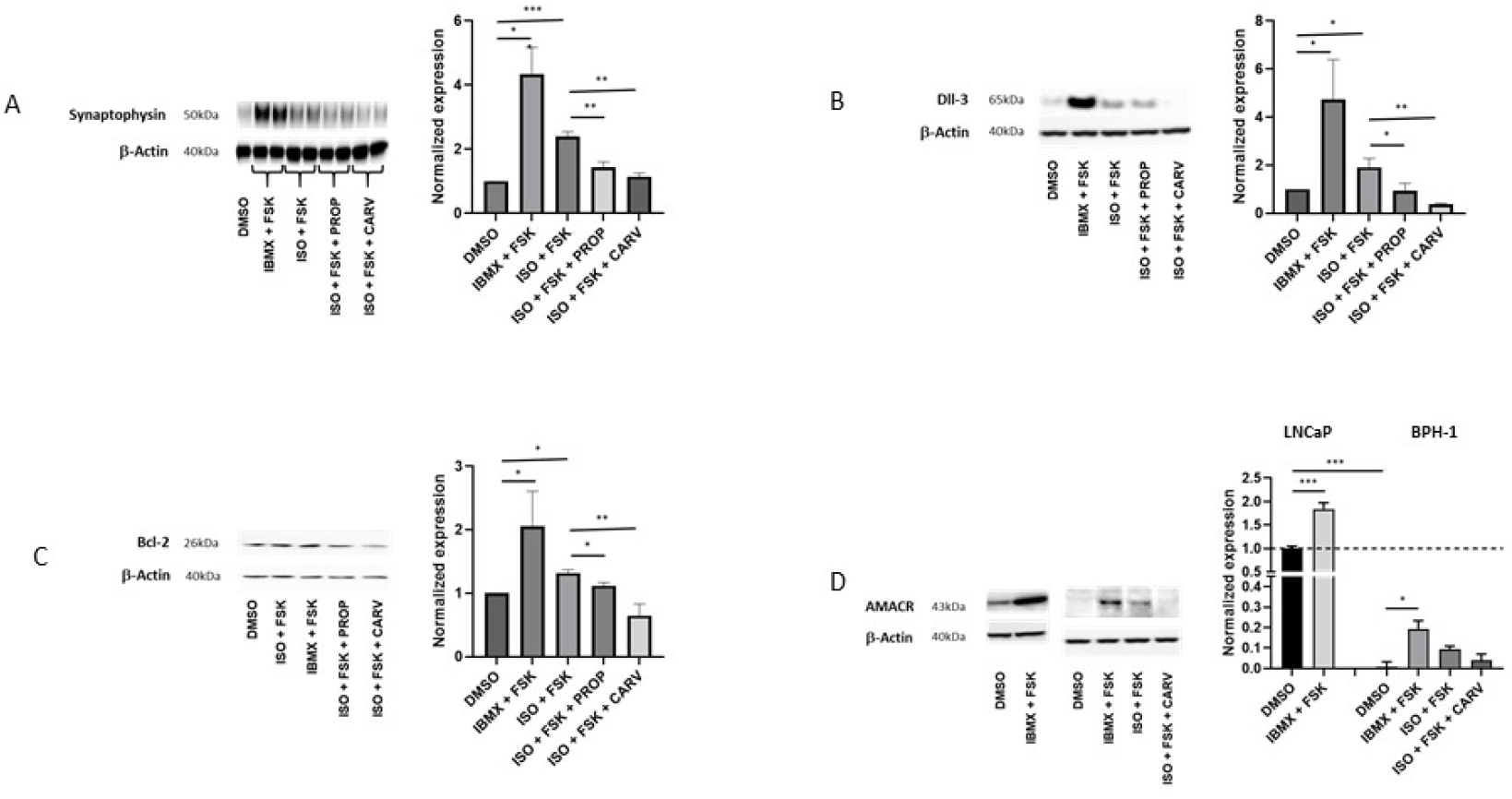
Activation of β adrenergic receptors induces neuroendocrine differentiation of BHP-1 cells. (A-C) Left: Western blot analyses for A) SYN, B) Dll-3, and C) Bcl-2 after the stimulation of BPH-1 cells for 10 days with 100 µM IBMX and 10 µM FSK or by 1 µM ISO and 10 µM FSK with or without 20 µM propranolol (PROP) or 5 µM carvedilol (CARV). Right: Quantification of western blot data (n=3-4). (D) Left: Western blot analyses for AMACR expression in LNCaP cells in the presence or absence of IBMX and FSK and in BPH-1 cells treated with IBMX and 10 µM FSK or ISO and FSK with or without CARV. Right: Quantification of western blot data (n=4).

**Figure 5:**
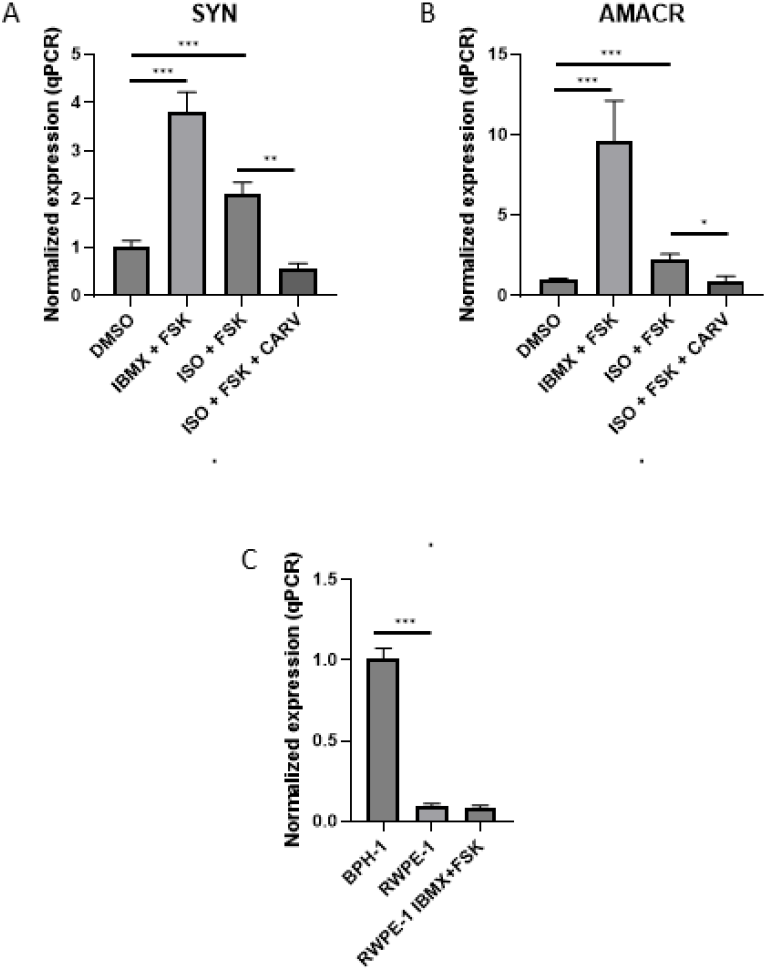
BPH-1 cells are more susceptible than normal cells to differentiation upon β2 adrenergic receptor stimulation. (A and B) RT-qPCR analyses of A) *SYN* mRNA (n=3-12) and B) *AMACR* mRNA (n=4-11) in BPH-1 cells treated for 10 days with 100 µM IBMX and 10 µM FSK or with 1 µM ISO and 10 µM FSK with or without 5 µM carvedilol (CARV). (C) RT-qPCR analyses of *SYN* mRNA in BPH-1 cells and in RWPE-1 cells treated or not with 100 µM IBMX and 10 µM FSK (n=6).

## Discussion

Here we provide the first evidence that NE-like cells can arise from a BPH model cell line. Although the presence of NE-like cells correlates with poor prognosis for prostate cancer patients, these cells are present not only during the later phase of this disease, as described before (32), but may also precede the occurrence of BPH in hypertensive patients (22). Previous work demonstrated a link between β2 adrenergic receptor expression and activity and the presence of NE-like cells during the late phase of prostate cancer (33), and our data suggest that β2 adrenergic receptor activation results in NED in the BPH transition zone. It is known that these β2 adrenergic receptors are present in prostate tissues, and their expression is correlated to the grade of prostate cancer (33). Indirect evidence of their activity comes from the well-known BPH treatment using α adrenergic receptor blockers (34). Considered together, these data suggest that there is a constitutive release of neurotransmitters in the prostate transition zone.

Blockers of β1 adrenergic receptors are preferentially used to treat hypertension, and their lack of effect in our study suggests that the use of β2 blockers to prevent NE-like cell appearance should be reconsidered. It is worth noting that the use of β2 blockers is beneficial in breast and lung cancers (35, 36). Beta blocker concentration must be adjusted to prevent the cytotoxic effects observed with propranolol and carvedilol in BPH-1 cells, which were previously described in other tissues (31). Although β2 blocker side effects are well documented (37), drug repurposing suggests that their use could benefit to BPH patients with adenocarcinoma as well as patients undergoing androgen deprivation therapy (38).

It has recently become clear that stress responses, which occur during the immediate post-operative period and which result in the release of catecholamines such as epinephrine and norepinephrine, could mediate many of the adverse effects of surgery through their direct effects on the malignant tissue and through modulation of host physiology and immune competence (1, 39). Even if BPH and cancer are two separate prostate diseases, the effects of β2 adrenergic receptor agonists on BPH-1 cells shown in our study suggest that a systematic search for NE-like cells in the transition zone in these patients after BPH surgery or radical prostatectomy in a large cohort should be conducted. If these cells are present in a large subset of patients, β2 blocker use should be recommended.

The two neurotransmitters epinephrine and norepinephrine have similar affinities for both α and β adrenergic receptors, and α blockers such as tamsulosin and silodosin are used to treat BPH as they induce smooth muscle relaxation (34). BPH-1 cell proliferation was not affected by phenylephrine, a specific α1 adrenergic receptor agonist, suggesting that development of NE-like cells results specifically from activation of β2 adrenergic receptors.

ISO concentration range (1-100µM) used in our study was similar to those previously described in prostate cancer cell lines (14, 40) but it is interesting to note that cAMP production induced by this β2 adrenergic receptor agonists occurs at submicromolar concentrations in the same cell types (28). It is quite possible that NED requires high and prolonged cAMP production to be established in time. Such high ISO concentrations are likely to be achieved *in vivo* as large neurotransmitter concentrations are often measured at the junction between the fibers and the target cells. For example, this kind of pattern (duration of the stimulation and high concentrations) has been described for glutamate metabotropic receptors (41).

Previous transcriptome profiling showed clear differences between normal and BPH tissues (42). As activating the cAMP/PKA pathway in RWPE-1 cells, a normal prostate tissue line, did not result in the activation of NE-like markers in our hands, we speculate that BPH tissues differ from healthy tissues in their response to β2 adrenergic receptor stimulation. The mechanism by which LNCaP and BPH-1 cells are transformed into NE-like cells is not fully understood, and it will be of interest to determine why RWPE-1 and BPH-1 cells respond differently to β2 adrenergic receptor agonists. AMACR expression was detected by qPCR and western blot in BPH-1 cells treated with β2 adrenergic receptor agonists and cAMP/PKA pathway activators for 10 days. The low level of AMACR expression in BPH-1 cells compared to LNaP cells suggests that additional triggers or a longer duration of stimulation may be needed to fully transform BPH-1 cells into NE-like cells. Very low levels of AMACR protein expression were observed in previous studies of RWPE-1 and BPH-1 cells (43).

CREB is a relay for the cAMP/PKA pathway that triggers NED (40). Androgen deprivation therapy and radiotherapy induce NED via the CREB pathway (44-46). Enzalutamide increases NED in LNCaP cells (47), and clinical studies have demonstrated that this drug increases the frequency of NE-like cells leading to poorer outcome for men with prostate cancer (48). In contrast, the use of antihypertensive medication significantly increased survival of patients with metastatic castration-resistant prostate cancer (49). NED is associated to a decrease in androgen receptor expression (15), and as all these treatments as well as stress leads to an increase in NE-like cells, this may explain why late prostate cancer becomes insensitive to androgen deprivation therapy (15, 50, 51). For patients with metastatic castration-resistant prostate cancer treated with abiraterone or docetaxel, NED and the proportion of NE-like cells were critical predictive factors, so NED might serve as a biomarker to predict outcomes and to guide treatment decisions (52).

## Conclusion

Stress is positively correlated with cancer initiation, progression, and metastasis in several types of cancer (53-55). Our data indicate that stress neurotransmitters such as epinephrine and norepinephrine trigger NED even in cells that serve as a model for BHP. This mechanism, which involve cAMP/PKA and CREB pathways, may underly stress-mediated induction of other types of cancer. Combining use of enzalutamide (and other drugs used to treat prostate cancer) and β2 blockers may reduce the appearance of NE-like cells, which are associated with poor outcome in this disease (56).

## Supporting information

**Supp. Figure 1:**

**Combination treatment with IBMX and FSK reduces BPH-1 cell proliferation**.

A) Left: Mean increase in BPH-1 cells after 96h cultured without (CTRL) and with 0.02% DMSO (n=30). Right: Plot of BPH-1 cell number increase after 96h revealing variability (range 4 to 28). (B) Left: Plot of normalized DMSO values to mean distilled water values versus time. Right: Plot of normalized control values revealing variability (range 0.78 to 1.24). (C) Left: Proliferation of RWPE-1, BPH-1, LNCaP, DU-145, and PC-3 cells at 96 h in the presence of IBMX and FSK normalized to proliferation in DMSO (n=3, 23, 16, 8, and 8, respectively). Right: Numbers of RWPE-1 cells (log scale) treated with DMSO (black squares) or the combination of IBMX and FSK (black circles) versus time in days (n=4).

**Supp. Figure 2:**

**β-adrenergic receptor agonists have little effect on viability of BPH-1 cells**.

Cell viability was assessed after 4 days in the presence of the IBMX and FSK (I+F), ISO), propranolol (PROP), or carvedilol (CARV) (n=6 to 8 for treatments and n=32 for untreated control cells).

**Supp. Figure 3:**

**The β-adrenergic receptor agonist isoproterenol reduces proliferation of prostate cancer lines**.

Normalized cell proliferation was assessed as a function of concentration of ISO for RWPE-1 (n=3-4), LNCaP (n=6-10), and PC-3 (n=4-5) prostate cell lines.

**Supp. Figure 4:**

**Activation of the β2-adrenergic receptor blocks BPH-1 cell proliferation**.

Cell proliferation measured HOW in the presence of epinephrine (EPI, n=4-7), norepinephrine (NOR, n=7-8), phenylephrine (PHE, n=3-4), and mirabegron (MBG, n=7).

**Supp. Figure 5:**

**Beta blockers inhibit prostate cancer cell proliferation**.

Normalized cell proliferation as measured HOW in RWPE-1 (n=3-4), LNCaP (n=4-8), and PC-3 (n=3-4) prostate cell lines in the presence of β2 blockers 20 µM propranolol (PROP) and 5 µM carvedilol (CARV) and 20 µM atenolol (ATE).

**Supp. Figure 6:**

**Uncropped western blots**

Uncropped western blots showing all the bands with all molecular weight markers for SYN (A), Dll-3 (B), Bcl-2 (C) and AMACR (D) in control conditions (DMSO) and in the presence of IBMX and FSK or ISO 1µM and FSK in the presence or absence of 20µM propranolol (PROP) or 5µM carvedilol (CARV).

